# Ancient evolution of hepadnaviral paleoviruses and their impact on host genomes

**DOI:** 10.1101/2020.11.02.364562

**Authors:** Spyros Lytras, Gloria Arriagada, Robert J. Gifford

**Author notes:** **Spyros Lytras:** *MRC-University of Glasgow Centre for Virus Research, 464 Bearsden Rd, Bearsden, Glasgow, UK, G61 1QH. **Gloria Arriagada:** *Instituto de Ciencias Biomedicas, Facultad de Medicina y Facultad de Ciencias de la Vida, Universidad Andres Bello, Echaurren 183, Santiago, Chile. **Robert J Gifford:** *MRC-University of Glasgow Centre for Virus Research, 464 Bearsden Rd, Bearsden, Glasgow, UK, G61 1QH. **Correspondence:** Robert J Gifford.

## Abstract

Hepadnaviruses (family *Hepadnaviviridae*) are reverse-transcribing animal viruses that infect vertebrates. Vertebrate genomes contain DNA sequences derived from ancient hepadnaviruses, and these ‘endogenous hepatitis B viruses’ (eHBVs) reveal aspects of the long-term coevolutionary relationship between hepadnaviruses and their vertebrate hosts. Here, we use a novel, data-oriented approach to recover and analyse the complete repertoire of eHBV elements in published animal genomes. We show that germline incorporation of hepadnaviruses is exclusive to a single vertebrate group (Sauria) and that the eHBVs contained in saurian genomes represent a far greater diversity of hepadnaviruses than previously recognised. Through in-depth characterisation of eHBV elements we establish the existence of four distinct subgroups within the genus *Avihepadnavirus* and trace their evolution through the Cenozoic Era. Furthermore, we provide a completely new perspective on hepadnavirus evolution by showing that the metahepadnaviruses (genus *Metahepadnavirus*) originated >300 million years ago in the Paleozoic Era, and has historically infected a broad range of vertebrates. We also show that eHBVs have been intra-genomically amplified in some saurian lineages, and that eHBVs located at approximately equivalent genomic loci have been acquired in entirely distinct germline integration events. These findings indicate that selective forces have favoured the accumulation of hepadnaviral sequences at specific loci in the saurian germline. Our investigation provides a range of new insights into the long-term evolutionary history of reverse-transcribing DNA viruses and demonstrates that germline incorporation of hepadnaviruses has played an important role in shaping the evolution of saurian genomes.

## BACKGROUND

Hepadnaviruses (family *Hepadnaviridae*) are reverse-transcribing DNA viruses that infect vertebrates. The type species - hepatitis B virus (HBV) - is estimated to infect ~300 million people worldwide, causing substantial morbidity and mortality. Hepadnaviruses have enveloped, spherical virions and a small, circular DNA genome ~3 kilobases (Kb) in length. The genome is characterised by a highly streamlined organization incorporating extensive gene overlap - the open reading frame (ORF) encoding the viral polymerase (P) protein occupies most of the genome and typically overlaps at least one of the ORFs encoding the core (C), and surface (S) proteins.

For decades only two hepadnavirus genera were known: genus *Orthohepadavirus*, which infects mammalian species, and genus *Avihepadnavirus*, which infects avian species. Since 2019, however, five hepadnavirus genera are recognised [1]. The three newly defined genera include the herpetohepadnaviruses (genus *Herpetohepadnavirus*), which infect amphibians and reptiles, as well as two highly distinct groups that infect fish - the metahepadnaviruses (genus *Metahepadnavirus*) and the parahepadnaviruses (genus *Parahepadnavirus*) [2–4]. Unexpectedly, phylogenetic analysis revealed that the metahepadnaviruses are more closely related to the mammalian orthohepadnaviruses than to other hepadnaviral lineages, leading to proposals that inter-class transmission of hepadnaviruses between fish and terrestrial vertebrates has occurred in the past [3, 5].

Whole genome sequencing has revealed the presence of DNA sequences derived from hepadnaviruses in some vertebrate genomes. These ‘endogenous hepatitis B viruses’ (eHBVs) are thought to have originated via ‘germline incorporation’ events in which hepadnavirus DNA sequences were integrated into chromosomal DNA of germline cells and subsequently inherited as novel host alleles. Most eHBV sequences that arise in this way will be quickly purged from the gene pool via drift and natural selection. Occasionally, however, some may persist long enough to become genetically fixed in the germline of ancestral species. Fixed eHBVs are expected to remain in the germline indefinitely unless removed by macrodeletion, but in the absence of selective pressure their sequences will gradually degrade via neutral mutation.

Analysis of eHBVs has proven immensely informative with respect to the long-term evolutionary history of the *Hepadnaviridae*. eHBV sequences are in some ways equivalent to hepadnavirus ‘fossils’ in that they provide a source of retrospective information about the distant ancestors of modern hepadnaviruses. Before ancient eHBV sequences provided a means of calibrating the timeline of hepdnavirus evolution, the family was thought to have originated within the past 100,000 years. However, the discovery of ancient eHBV sequences exhibiting remarkable similarity to contemporary strains demonstrates that hepadnaviruses infected vertebrate ancestors millions of years ago, during the Mesozoic and Cenozoic Eras [6–9]. All eHBVs identified so far derive from viruses belonging to the *Avihepadnavirus* or *Herpetohepadnavirus* genera.

Currently, the distribution and diversity of hepadnavirus-related sequences in animal genomes remains incompletely characterized. Studies have shown that multiple additional, lineage-specific eHBV insertions are present in some species [7, 10, 11]. However, progress in characterising these elements has been hampered by the challenges inherent in analysing large numbers of fragmentary and degenerated eHBV sequences. In this investigation we sought to directly address these challenges and comprehensively map the distribution and diversity of eHBV sequences in vertebrate genomes. Through comparative and phylogenetic analysis of the eHBV sequences identified in our study, we derive a wide range of novel insights into the evolution of hepadnaviruses and their impact on animal genomes.

## METHODS

### Genome screening in silico

We used the database-integrated genome screening (DIGS) tool [12] to derive a non-redundant database of loci within published WGS assemblies that show similarity to hepadnavirus-specific polypeptides. The DIGS tool is written using the PERL scripting language and implements a ‘database-integrated’ genome screening framework by using the MySQL relational database management system (RDBMS) to: (i) coordinate systematic screening and; (ii) capture output data. Similarity searches are performed using the basic local alignment search tool (BLAST) program suite [13]. The framework requires the collation of a reference sequence library. This provides a source of ‘probes’ (for searching WGS data using the tBLASTn program). Additionally, the set of sequences that is recovered via BLAST-based screening can be classified via comparison to curated set of reference sequences (this time using the tBLASTx program). For this project, we collated a library comprised of genome length sequences of representative hepadnavirus species (**Table S1**) and previously characterised eHBVs (see **Table S2**). In addition, we included the sequences of retroelements disclosing similarity to hepadnaviruses, which could be expected to produce false positive matches to hepadnavirus probes [14]. Whole genome sequence data were obtained from the National Center for Biotechnology Information (NCBI) genome database resource. We obtained all vertebrate genomes available as of March 2020.

For all five hepadnavirus genera we generated a multiple sequence alignment (MSA) that contained full-length sequences of all genus members (i.e. virus species and distinct eHBV insertions). MSAs were generated using a combination of MUSCLE and a BLAST-based, codon aware alignment method implemented in GLUE [15]. Genome length alignments were manually inspected and adjusted to correct problematic regions associated with germline mutations and insertions in eHBVs. The reference library was used to derive a set of polypeptide sequences as probes and references in DIGS (**Table S1**). For eHBVs we translated putative ORFs (inferred during the alignment process described above) to obtain representative polypeptide sequences. Sequences disclosing similarity to hepadnaviral probes were classified via tBLASTx-based comparison to this sequence set.

Via DIGS we generated a database of genomic sequences disclosing similarity to hepadnaviruses. We extended the core schema of this database to incorporate additional tables representing the taxonomic classifications of viruses, eHBVs and host species included in our study. We used structured query language (SQL) to interrogate this database, filtering sequences based on their similarity to reference sequences, the taxonomic properties of the closest related reference sequence, and the taxonomic distribution of related sequences across hosts. Using this approach we categorised sequences into: (i) putatively novel eHBV elements; (ii) orthologs of previously characterised eHBVs (e.g. copies containing large indels); (iii) non-viral sequences that cross-matched to hepadnavirus probes (e.g. retrotransposons). Sequences that did not match to previously reported eHBVs were further investigated by incorporating them into our genus-level, genome-length MSA along with all of our reference taxa and reconstructing maximum likelihood phylogenies using RAxML (version 8) [16].

Where phylogenetic analysis supported the existence of a novel eHBV insertion, we also attempted to: (i) determine its genomic location relative to annotated genes in reference genomes; and (ii) identify and align eHBV-host genome junctions and pre-integration insertion sites (see below). Where these investigations revealed new information (e.g. by confirming the presence of a previously uncharacterised eHBV insertion) we updated our reference library accordingly. This in turn allowed us to reclassify all of the putative eHBV loci in our database and group sequences more accurately into categories. By iterating this procedure we progressively resolved the majority of eHBV sequences identified in our screen into groups of orthologous sequences derived from the same initial germline incorporation event (**Table S3**). eHBV elements were given unique IDs using a systematic approach, following a convention established for endogenous retroviruses [17].

### Comparative analysis of hepadnavirus and eHBV sequences

We used the GLUE software environment [15] to create a sequence data resource (‘Hepadnaviridae-GLUE’) capable of supporting reproducible comparative investigations of hepadnavirus genomes - including those that utilise host genomic data such as eHBVs (**Fig. 1a**). We created a GLUE project that contains all of the data items associated with our investigation (i.e. virus genome sequences, multiple sequence alignments, genome feature annotations, and other sequence-associated data) and uses a relational database to represent the semantic relationships between them. Representative genome sequences for hepadnavirus species recognised by ICTV were obtained from GenBank. Sequences of recently described hepadnaviruses not yet available in GenBank were obtained from study authors [4].

**Figure 1.**
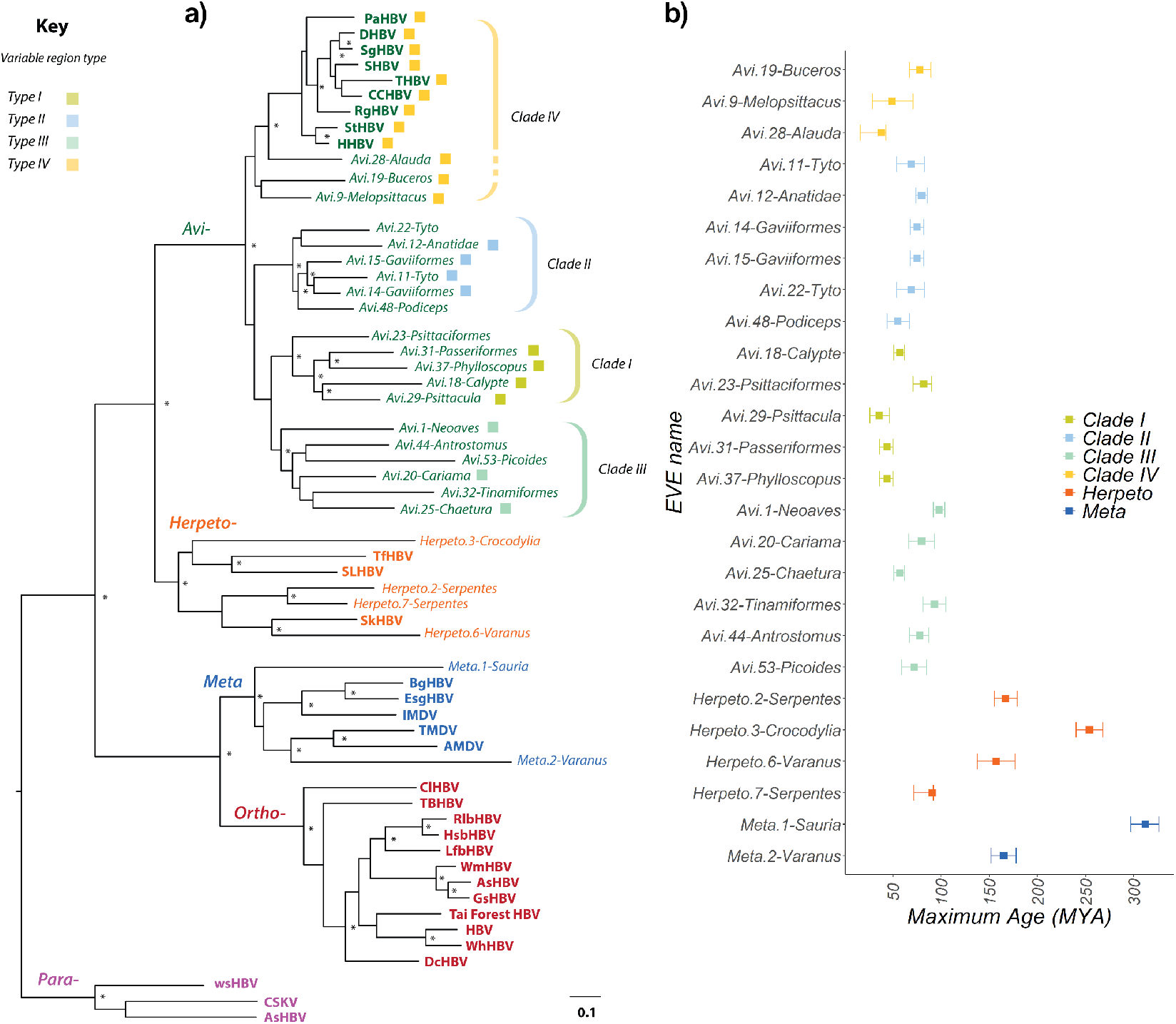
Recovery of paleovirus sequences reveals the evolutionary history of hepadnaviruses. Panel **(a)** shows a maximum likelihood phylogeny constructed using codons 355-440 and 500-781 of the polymerase (P) protein (Hepatitis B virus coordinates, GenBank reference sequence accession number: NC_003977). The phylogeny is rooted on the Parahepadnavirus genus, based on the basal position of this genus in trees constructed using the Nackednaviruses as an outgroup (**Fig. S1**). Virus names are shown in bold. IDs of endogenous hepatitis B viruses (eHBVs) are shown in italic. Taxon label colours correspond to viral genera as follows: purple=Parahepadnavirus; blue=Metahepadnavirus; red=Orthohepadnavirus; orange=Herpetohepadnavirus; green=Avihepadnavirus. Virus name abbreviations are as shown in **Table S1**. Brackets to the right indicate subclades within the avihepadnavirus genus. The dashed bracket for Clade IV denotes that grouping of eHBV elements 9, 19 and 28 does not have high support here, but is supported by other phylogenetic evidence. Coloured squares next to avihepadnavirus taxon labels variable region ‘type’ as indicated by the key. Some eHBV sequences do not span this region and thus cannot be assigned. Asterisks indicate nodes with bootstrap support >=70, based on 1000 replicates. The scale bar shows evolutionary distance in substitutions per site. The plot in panel **(b)** shows the window in geological time during which we estimated various eHBV insertions to have been generated, based on their distribution across vertebrate taxa. Abbreviations: MYA=Million years ago.

Hepadnavirus sequences were virtually ‘rotated’ within GLUE as required to represent them within the same coordinate space (i.e. using the same genomic start position). Using GLUE we implemented an automated process for constructing alignments of putatively orthologous sequences and thereafter using these alignments to derive consensus sequences representing each set of orthologs. Once the presence of a novel eHBV insertion was established, a consensus sequence representing this insertion was incorporated into the appropriate genome-length MSAs for the genus it derived from (this could usually be inferred via sequence similarity, but in marginal cases was confirmed by phylogenetic analysis using a broader taxa set).

Because all aligned eHBV and virus sequence data held in our project have been adjusted to occupy a standardised coordinate space (by default determined by our project master reference, hepatitis B virus), we could use functions implemented within the GLUE software to infer coverage relative to genus master reference genomes, and thereby infer the genomic structure of consensus eHBV elements. In addition, all the alignments constructed in our study were rationally linked to one another via a ‘constrained alignment tree’ data structure that links multiple sequence alignments (MSAs) constructed at distinct taxonomic levels. This allowed us to automate the reconstruction of evolutionary relationships between hepadnaviruses and eHBV elements at different taxonomic levels (e.g. family, genus, ortholog), and using the standardised coordinate space to select alignment partitions corresponding to specific genome features and subdomains.

### Genomic analysis

To confirm that the eHBV elements identified in our study were distinct from those previously reported (i.e. they derive from a distinct germline incorporation event) we investigated the locus surrounding each putatively novel eHBV. To identify flanking genes, we extracted 2kb sequences flanking each eHBV hits (using utility scripts implemented within the DIGS tool). We used BLASTn to identify the corresponding region in related species (i.e. a region that disclosed the expected degree of homology to both the upstream and downstream flanking regions). By viewing the region in the ENSEMBL genome browser, we obtained the unique identifiers of the most closely located genes in the regions upstream and downstream of eHBV insertions.

To assess potential presence of transposable elements (TE) around eHBVs of interest (**Fig 3a**) we extracted the 5kb sequences flanking the eHBV coordinates from the respective genome assemblies, adjusting for reverse complementarity. These sequences were analysed for TE presence using HMMER [28] against the Dfam HMM profile library [29].

### Insertion dating

Dating of eHBV insertions presented in this study have been estimated by examining the most distant host species sharing a particular eHBV and using these hosts’ divergence date and confidence intervals (CI) as reported in Timetree (ref: www.doi.org/10.1093/molbev/msx116).

## RESULTS

### Endogenisation of hepadnaviruses is unique to Saurian species

We screened all available WGS data for metazoan species and identified >930 sequences disclosing similarity to hepadnaviruses (**Table 1**, **Table S3**). We found that *bona fide* eHBV elements are only present in the genomes of saurian species. Furthermore, reconstruction of the phylogenetic relationships between eHBVs and contemporary hepadnaviruses revealed that saurian genomes contain a broader diversity of eHBV elements than previously recognised, with elements derived from the metahepadnavirus-like elements being present, as well as elements derived from the *Avihepadnavirus* and *Herpetohepadnavirus* genera (**Fig. 1a, Fig. S1a**).

**Table 1.** EHBV loci detected in vertebrate genomes.

| eHBV element ID <sup>a</sup> | Virus Clade<sup>b</sup> | Num. Species<sup>c</sup> | Num. Seqs<sup>d</sup> | Flanking genes<sup>e</sup> |  | Min age <sup>f</sup> | Max age <sup>g</sup> |
| --- | --- | --- | --- | --- | --- | --- | --- |
|  |  |  |  | Upstream | Downstream |  |  |
| **Metahepadnavirus** |  |  |  |  |  |  |  |
| Meta.1-Sauria |  | 27 | 27 | *KLF8* | *ENSAC* | 280 (273-286) | 312 (297-326) |
| Meta.2-Varanus |  | 1 | 1 | *NPVF* | *NFE2L3* | - | 165 (152-178) |
| Meta.3-Paroedura |  | 1 | 1 | *n/k* | *n/k* | - | 96 (83-98) |
| Meta.4-Pelusios |  | 1 | 1 | *LCP1* | *RUBCNL* | - | 99 (70-128) |
| Meta.5-Pelusios |  | 1 | 1 | *KCNA5* | *NTF3* | - | 99 (70-128) |
| Meta.6-Sphenodon |  | 1 | 1 | *n/k* | *n/k* | - | 252 (241-263) |
| **Herpetohepadnavirus** |  |  |  |  |  |  |  |
| Herpeto.1-Serpentes* | Snake | 9 | 9 | *TSHZ1* | *ZNF516* | 62 (49-74) | 91 (72-92) |
| Herpeto.2-Serpentes* | Snake | 10 | 10 | *NPFFR2* | *FTH1* | 62 (49-74) | 167 (155-179) |
| Herpeto.7-Serpentes | Snake | 3 | 3 | *ATP2A3* | *ZZEF1* | 9.2 (5.8-18.8) | 91 (72-92) |
| Herpeto.8-Serpentes | Snake | 5 | 5 | *OBI1* | *RBM26* | 62 (49-74) | 91 (72-92) |
| Herpeto.6-Varanus | Lizard | 1 | 1 | **WSCD2** | **WSCD2** | - | 157 (138-177) |
| Herpeto.3-Crocodylia* | Croc. | 1 | 1 | *NUP210* | *IQSEC1* | 44 (25-64) | 254 (240-268) |
| Herpeto.4-Crocodylia* | Croc. | 1 | 1 | *SORT1* | *PPIL1* | 26.7 (22-29) | 254 (240-268) |
| Herpeto.5-Testudines* | Turtle | 18 | 18 | *GBE1* | *ROBO1* | 184 (161-206) | 254 (240-268) |
| **Avihepdnavirus** |  |  |  |  |  |  |  |
| Avi.18-Calypte | I | 1 | 1 | **ENSSCUG00000017518** | **ENSSCUG00000017518** | - | 57 (51-62) |
| Avi.23-Psittaciformes | I | 11 | 71 | *n/a* | *n/a* | 49 (29-71) | 82 (71-90) |
| Avi.29-Psittacula | I | 1 | 1 | **KIDINS220** | **KIDINS220** | - | 36 (26-46) |
| Avi.31-Passeriformes† | I | 7 | 7 | *TIMM21* | *NETO1* | 38 (16-43) | 44 (36-50) |
| Avi.37-Phylloscopus | I | 2 | 2 | *EPHA3* | *ZNF654* | 1.2 | 44 (36-50) |
| Avi.49-Psittaciformes | I | 14 | 14 | **GRID2** | **GRID2** | 38 (30-50) | 82 (71-90) |
| Avi.52-Melopsittacus | I | 1 | 1 | *LUZP2* | *ANO3* | - | 38 (14-55) |
| Avi.11-Tyto | II | 1 | 1 | *NAV3* | *E2F7* | - | 69 (54-83) |
| Avi.12-Anatidae | II | 5 | 9 | **CCDC58** | **CCDC58** | 30 (26-35) | 80 (74-86) |
| Avi.14-Gavia | II | 1 | 1 | *TAS1R3* | *DVL1* | - | 75 (68-82) |
| Avi.15-Gavia | II | 1 | 1 | **TMEM182** | **TMEM182** | - | 75 (68-82) |
| Avi.22-Tyto | II | 1 | 1 | *MYBL2* | *PTPR* | - | 69 (54-83) |
| Avi.24-Apaloderma | II | 1 | 1 | **CDH23** | **CDH23** | - | 72 (59-85) |
| Avi.27-Sulidae | II | 4 | 233 | *n/a* | *n/a* | 9.6 | 73 (59-87) |
| Avi.35-Calypte | II | 1 | 1 | *CYB5A* | *TIMM21* | - | 57 (51-62) |
| Avi.46-Psittaciformes | II | 10 | 10 | *FMN1* | *RYR3* | 49 (29-71) | 82 (71-90) |
| Avi.48-Podiceps | II | 1 | 1 | *MATN1* | *PTPRU* | - | 55 (44-67) |
| Avi.1-Neoaves* | III | 110 | 111 | **FRY** | **FRY** | 85 (77-94) | 98 (92-104) |
| Avi.20-Cariama | III | 1 | 1 | *THBS1* | *KATNBL1* | - | 80 (66-93) |
| Avi.25-Chaetura | III | 1 | 1 | *KCNV1* | **ENSTGUG00000027711** | - | 57 (51-62) |
| Avi.32-Tinamiformes | III | 2 | 2 | *OLFM4* | *n/k* | 49 (37-62) | 93 (81-105) |
| Avi.34-Leptosomus | III | 1 | 1 | **LRR7** | **LRR7** | - | 70 (58-82) |
| Avi.42-Passeriformes† | III | 5 | 5 | **HECA** | **HECA** | 48 (33-50) | 82 (75-90) |
| Avi.44-Anstrostromus | III | 1 | 1 | *KCNV1* | *CSMD3* | - | 78 (67-87) |
| Avi.53-Picoides | III | 1 | 1 | *TRIB2* | *LPIN1* | - | 72 (59-85) |
| Avi.28-Alauda | IV | 1 | 1 | *EPHA6* | *NSUN3* | - | 38 (16-43) |
| Avi.4-Passeriformes†* | IV | 9 | 9 | **CDH23** | **CDH23** | 38 (16-43) | 82 (75-90) |
| Avi.5-Passeriformes†* | IV | 5 | 5 | *LMO3* | *MGST1* | 38 (16-43) | 82 (75-90) |
| Avi.6-Estrildinae* | IV | 2 | 2 | *FOXO3* | *ATG4C* | 10.1(8.7 - 11.6) | 11.8 (11-14) |
| Avi.8-Australiaves* | IV | 18 | 28 | **ATP2B2** | **ATP2B2** | - | 82 (71-90) |
| Avi.38-Passeriformes | IV | 8 | 8 | *TGIF2* | *AAR2* | 38 (16-43) | 82 (75-90) |
| Avi.9-Melopsittacus* | IV | 1 | 1 | **CD109** | **CD109** | - | 49 (29-71) |
| Avi.19-Buceros | IV | 1 | 1 | *PCDH18* | *PCDH10* | - | 78 (67-89) |
| Avi.7-Passeriformes†* |  | 6 | 6 | *TMEM132E* | *LIG3* | 38 (16-43) | 82 (75-90) |
| Avi.13-Paleognathia |  | 8 | 8 | **ENSDNVG00000017897** | *BCKDHB* | 93 (81-105) | 111 (105-118) |
| Avi.16-Turaco |  | 1 | 1 | *TMEM8B* | **ENSACUG00000013925** | - | 74 (63-85) |
| Avi.21-Paleognathia |  | 8 | 8 | *LMCD1* | *GRM7* | 93 (81-105) | 111 (105-118) |
| Avi.26-Psittaciformes |  | 7 | 8 | **NELL1** | **NELL1** | 38 (14-55) | 82 (71-90) |
| Avi.30-Anatidae |  | 5 | 5 | **ENSACDG00005009727** | *CCDC58* | 30 (26-35) | 80 (74-86) |
| Avi.39-Passeriformes† |  | 9 | 9 | **FXN** | **FXN** | 38 (30-50) | 82 (75-90) |
| Avi.41-Psittaciformes |  | 4 | 4 | *PHAX* | *KLF4* | 38 (30-50) | 82 (71-90) |
| Avi.43-Gallirallus |  | 1 | 1 | **DEND4A** | **DEND4A** | - | 64 (52-75) |
| Avi.45-Psittaciformes |  | 8 | 8 | **RNF38** | **RNF38** | 38 (30-50) | 82 (71-90) |

**Footnote:** <sup>a</sup> eHBV elements were given unique IDs using a systematic approach, following a convention developed for endogenous retroviruses [17]. Each is assigned a unique identifier (ID) constructed from three components. The first component (not shown here) is the classifier 'eHBV' denoting an endogenous hepadnaviral element. The second component comprises: (i) the name of the hepadnavirus genus the element derived from and; (ii) a numeric ID that uniquely identifies a specific integration locus, or for multicopy lineages, a unique founding event. The final component denotes the taxonomic distribution of the element. \* Previously reported elements eHBVs distributed across unranked clades. <sup>b</sup> Virus taxonomic groups below genera level are shown where known. <sup>c</sup> Number of species in which the insertion/lineage was identified. <sup>d</sup> Total number of orthologs/duplicated identified in screen. <sup>e</sup> Names of annotated genes flanking each insertion. Intronic insertions are shown in bold. EMSEMBL gene IDs are shown for genes that do not yet have names. n/k = not known; n/a = not applicable to multi-copy lineages; <sup>f</sup> Minimum age as determined via orthology and based on divergence times provided by TimeTree [19]; <sup>g</sup> Maximum age based on presence of empty insertion site in a sister clade, or other evidence (see Methods).

Relatively large numbers of avihepadnavirus-derived eHBVs were identified in avian genomes (**Table 1**), most of which represent only short, sub-genomic fragments. However, we identified 17 that represented complete, or near complete viral genomes (**Fig. 2**). We also identified previously unreported, herpetohepadnavirus-derived eHBVs in the genomes of a lizard (superorder Lepidosauria) and in snakes (order Serpentes). The novel snake elements were closely related to those previously reported in snake genomes [7] (**Fig 1a**, **Fig S1**), while the lizard element was identified in the Komodo dragon (*Varanus komodoensis*). *Herpeto.6-Varanus* was found to cluster robustly with skink hepatitis B virus (SkHBV) [4].

**Figure 2.**
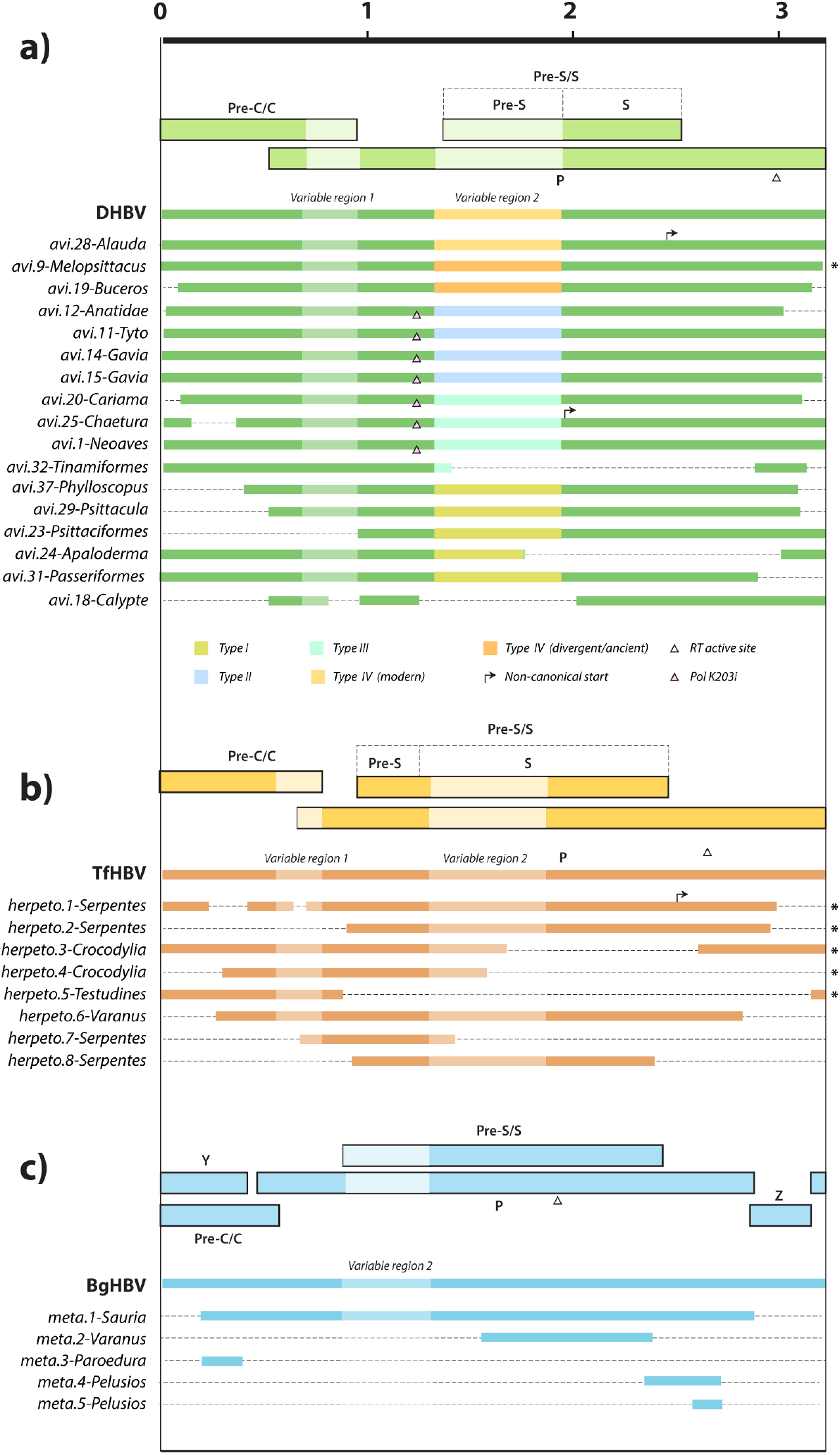
Genomic organisation of selected endogenous hepadnaviral element (eHBV) sequences identified in this study. eHBV structures are shown relative to the genomes of prototype virus species from the corresponding genus. Virus names are shown in bold, eHBV names are shown in italic. Thinner bars represent nucleic acid sequences. Thicker bars represent open reading frames in viral sequence. Asterisks indicate sequences that have been reported previously. Scale bar indicates sequence length in kilobases. Key shows relationships between symbols/shading and genome features. Abbreviations: DHBV (duck hepadnavirus); TfHBV (Tibetan frog hepadnavirus); BgHBV (bluegill hepadnavirus); Pre-C/C (Pre-Core/Core); P (Polymerase); Pre-S/S (Pre-Surface /Surface);

**Figure 3.**
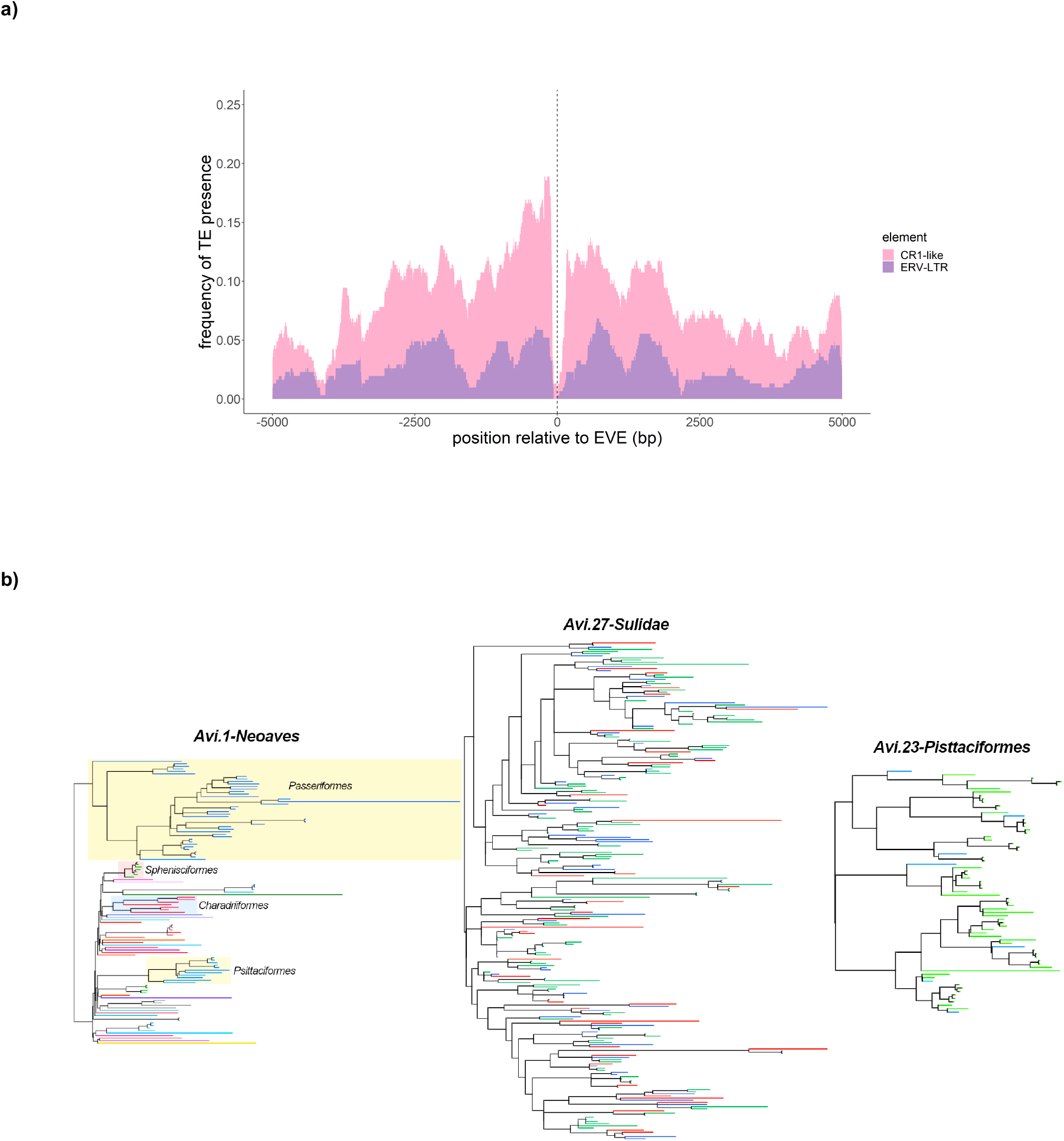
Genomic characteristics of multicopy eHBV lineages. **(a)** The plot shows the frequency with which specific transposable element (TE) sequences were detected in 5kb regions flanking 307 distinct members of the multicopy eHBV lineage *Avi.27-Sulidae*. Sequences were analysed for the presence of transposable elements (TE) using HMMER [30] against the Dfam HMM profile library [31]. Based on their descriptions, TEs detected in flanking sequences were divided into two categories: (i) related to the chicken repeat 1 group of retrotransposons (CR1) (shown in pink); (ii) related to endogenous retrovirus (ERV) long terminal repeat (LTR) (shown in purple); **(b)**Phylogenies of multicopy element lineages. (i) *Avi.1.Neoaves*; (ii) *Avi.27.Sulidae*; (iii) *Avi.23.Psittaciformes*. The terminal branches of all tips in the phylogenies are coloured based on their hosts’ taxonomic classification. The *Avi.1.Neoaves* phylogeny is labelled on the order level, *Avi.27.Sulidae* on the genus level and *Avi.23.Psittaciformes* on the family level. The order names of clear clades with multiple representatives are annotated on the *Avi1.Neoaves* phylogeny.

Notably, metahepadnavirus-like eHBV elements were identified in a wide range of saurian species, including birds, turtles, a lizard - the ocelot gecko (*Paroedura pictus*) - and the tuatara (*Sphenodon punctatus*). By contrast, herpetohepadnavirus-derived elements were only identified in reptiles, and avihepadnavirus-derived elements were only detected in birds.

### Several distinct avihepadnavirus lineages have circulated among birds during their evolution

Phylogenetic reconstructions demonstrate the presence of at least four distinct clades (I-IV) within the *Avihepadnavirus* genus (**Fig. 1**). Clade IV contains a mixture of extant avihepadnaviruses and eHBV insertions, while the remaining three clades are comprised exclusively of eHBV sequences. Notably, all four clades are highly divergent from one another in ‘variable region 2’ (which spans most of the Pre-S protein and includes regions that encode receptor-binding functions [18]), but within each clade these regions are relatively well conserved (**Fig. 2**, **Fig. S2a**). The order of ancestral branching among *Avihepadnavirus* clades is unclear – in phylogenies constructed using highly conserved regions of the P gene and rooted on herpetohepadnaviruses, none is clearly basal or derived relative to the others (data not shown). Notably, however, clade IV and clade II share a conserved, synapomorphic character: the insertion of a valine (V) or isoleucine (I) residue in the P protein, between positions 203 and 204 (**Fig. 2**, **Fig. S2b**). This shared, conserved character indicates that these two clades are more closely related to one another than they are to other hepadnaviruses - at least in the region around the synapomorphy. Overall, germline incorporation events involving each of the four avihepadnavirus clades seem to have occurred throughout the evolution of birds, with some occurring prior to major divergences in the avian tree, and others being confined to specific avian species or subgroups (**Table 1, Fig 1b**).

Near-complete insertions derived from clade I were identified in rose-necked parakeets (*Avi.29-Psittacula*) and in Anna’s hummingbird (*Calypte anna*) as well as two distinct elements in songbirds (order Passeriformes). Among the two songbird elements, one was found only in warblers (*Avi.37-Phylloscopus*) while another (*Avi.37-Passeriformes*) was found in five distinct families within the superfamily Passeroidea, establishing that it integrated into the passeroid germline >38 Mya (CI: 16-43 Mya). We also identified clade I-derived elements in parrots that represent only fragments of a hepadnavirus genome. These elements, which appear to have been intragenomically amplified (discussed below) and include some elements that are orthologous across all parrots (order Psittaciformes), indicate that germline incorporation occurred >49 Mya (CI: 29-71 Mya) prior to the divergence of the kea (*Nestor notabilis*) from other parrot lineages [19].

Clade II contains sequences derived from ducks (family Anatidae), red-throated divers (*Gavia stellata*) and barn owls (*Tyto alba*). The insertion in ducks was incorporated >30 Mya (CI 26-35 Mya), prior to the divergence of mallards (*Anas platyrhynchos*) and ruddy ducks (*Oxyura jamaicensis*). Notably, multiple, genome-length eHBV elements derived from this lineage were often identified in the same species or species group. For example, multiple, clade II-derived eHBVs were identified in both the *Tyto* (*Avi.11* and *Avi.22*) and *Gavia* (*Avi.14* and *Avi15*) germlines. However, in-depth analysis of these sequences shows that each derives from distinct germline incorporation events. Not only are they located in entirely distinct genomic loci (**Table 1**), they show higher divergence in the variable regions of their genome than in other more conserved regions (**Fig. S3**) – this is consistent with them being separated by multiple rounds of viral replication, rather than neutral divergence following an intragenomic duplication process. Notably, the greatest extent of divergence was observed in the regions of the genome that encode receptor binding functions.

Clade III includes the ‘*Avi.1-Neoaves*’ element (previous names include eAHBV-FRY [4] and eZHBVc [7]), which is the first avihepadnavirus-derived eHBV element to be reported, and is also the oldest. It is orthologous across the Neoaves clade, which includes all avian species except the paleognathes (infraclass Paleognathae) and fowl (Galloanserae; ducks, chickens, and allies). We identified additional eHBVs derived from this lineage in a broad range of avian groups. Notably, clade I-derived insertions are present in the paleognathe germline: the genomes of white-throated and Chilean tinamous contain orthologous eHBVs demonstrating that clade I avihepadnaviruses circulated in paleognathe birds >49 Mya (CI 37-62 Mya). In addition, we identified clade I-derived insertions in order Trogoniformes represented by the bar-tailed trogon (bar-tailed trogon), in clade Strisores, represented by the swift (*Chaetura pelagica*), and in clade Australiaves, represented by the red-legged seriema (*Cariama cristata*). This broad distribution is consistent with the demonstrably ancient origins of this lineage.

All clade IV-derived eHBVs group basal to the exogenous avihepadnaviruses, which cluster together as a derived, crown group within this clade. We identified a full-length insertion in the Eurasian skylark (*Alauda arvensis*) genome that shows a higher level of relatedness to modern hepadnaviruses than does any previously reported eHBV (**Fig 1a**). Notably, *eHBV-Avi.28-Alauda* was the only avihepadnavirus-derived eHBV element found to exhibit similarity to modern avihepadnaviruses in the variable region of the genome (**Fig. 2, Fig. S2a**). Some phylogenetic trees support the inclusion of eHBV elements previously reported in the budgerigar genome [11], and a newly identified element identified in the genome of the rhinoceros hornbill (*Buceros rhinoceros*), within clade IV (**Fig. S1g**).

### Avian genomes contain multicopy eHBV lineages

In addition to genome-length sequences, avian genomes contain multiple eHBV elements that represent only fragments of an avihepadnaviral genome. Furthermore, some avian lineages contain expanded sets of highly related eHBVs. Most strikingly, we identified >300 copies of a highly duplicated eHBV element in cormorants and shags (order Suliformes). This lineage, named *Avi.27-Sulidae*, appears to be derived from a single germline incorporation event involving an ancient, clade II avihepadnavirus (**Fig. S1g**), and is comprised of fragments spanning a short region at the 3’ terminal end of the *pol* gene. Investigation of *Avi.27* elements revealed that the vast majority are flanked on both sides by transposable element (TE) sequences (**Fig. 3a**), suggesting this multicopy lineage may have arisen in association with TE activity (i.e. integration into a TE led to an eHBV-derived sequence being mobilised). Phylogenies indicate that the initial germline incorporation event that gave rise to this multicopy eHBV lineage predates the diversification of the four cormorant species in which it was identified, as evidenced by the presence of multiple, multi-species sub-clusters in phylogenies (see **Fig. 3b**) and the presence of multiple orthologous integration sites (data not shown).

We also identified apparently intragenomically amplified, avihepadnavirus-derived eHBV elements in the genomes of parrots. In this case, the amplified elements appear to derive from an ancient clade I avihepadnavirus. Although the elevated eHBV copy number found in certain avian orders reflects intragenomic amplification, it is nonetheless clear that the rate of germline incorporation is significantly higher in birds than in any other vertebrate group. We characterised eHBV loci in saurian genomes by identifying the nearest annotated genes upstream and downstream of EVE integration sites (**Table 1)**. Excluding integrations that occurred as a result of intragenomic amplification, we estimate that at least 57 distinct germline incorporation events-each involving a distinct hepadnavirus progenitor - have occurred during avian evolution.

### eHBV elements are enriched at specific loci in the saurian germline

Strikingly, our analysis of genes flanking eHBV insertions identified several pairs of elements that are fixed at distinct, but nonetheless approximately equivalent genomic sites. We identified six cases in which eHBV elements that appear to derive from distinct germline incorporation events have been fixed at approximately equivalent genomic loci (**Table 2**). Almost all involve avihepadnaviruses independently integrating at similar loci in avian genomes. However, in most of these cases the two eHBV elements involved each derive from distinct clades within the *Avihepadnavirus* genus. Furthermore, we also identified one case in which a herpetohepadnavirus-derived element in snakes (*Herpeto.7*) is located at the same approximate position as a member of the avihepadnavirus-derived *Avi.23* lineage (**Fig 4a**).

**Table 2.** Pairs of apparently distinct EHBV insertions at adjacent loci.

| Pair name | First member |  | Second member |  |
| --- | --- | --- | --- | --- |
|  | Name | Genus (Clade)<sup>a</sup> | Name | Genus (Clade)<sup>a</sup> |
| 1 (ZZEF) | Avi.23-Psittaciformes | Avi- (I) | Herpeto.7-Serpentes | Herpeto- (Snake) |
| 2 (ANO5) | Avi.52-Melopsittacus | Avi- (I) | Avi.26-Psittaciformes | Avi- (NK) |
| 3 (CDH23 intronic) | Avi.20-Cariama | Avi- (III) | Avi.46-Psittaciformes | Avi- (II) |
| 4 (CYB5A & TIMM21) | Avi.35-Calypste | Avi- (II) | Avi.31-Passeriformes | Avi- (I) |
| 5 (FBXO15-NETO1) | Avi.24-Apaloderma | Avi- (II) | Avi.4-Passeriformes | Avi- (IV/V) |
| 6 (CCDC58 intronic) | Avi.12-Anatidae | Avi- (I) | Avi.30-Anatidae | Avi- (NK) |

Footnote: <sup>a</sup> Avi- = *Avihepadnavirus*; Herpeto- = *Herpetohepadnavirus*;

**Figure 4.**
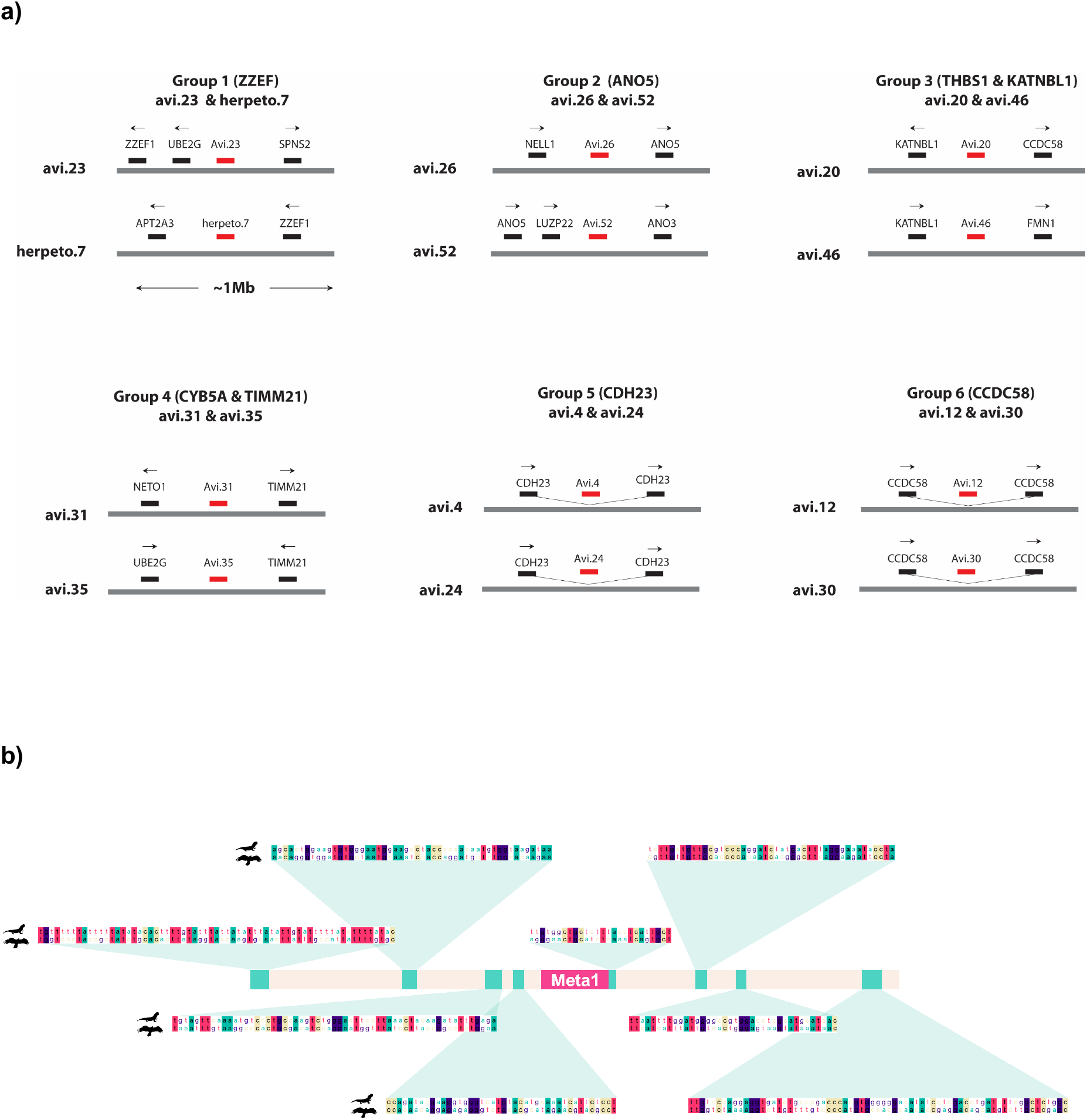
**(a)** Loci containing multiple fixed eHBV elements. Schematic representation six genomic loci where eHBV lineages originating in independent germline colonisation events have been fixed at adjacent positions. Grey bars represent genomic DNA. Black bars represent genes or exons with arrows showing the direction of transcription. Red bars represent eHBV elements. **(b)** Schematic representation of the *Sphenodon punctatus* copy of the *Meta.1-Sauropsida* (Meta1) genomic region including 1kb flanking regions to each side of the EVE. The sequence homology between the S. punctatus (top) and the *A. chrysaetos canadensis* (bottom) genome is highlighted in representative subregions around the orthologous EVE, labelled in green.

### Metahepadnaviruses circulated in the late Paleozoic Era

Most of the metahepadnavirus-like elements identified in our screen were comprised of short fragments ~300-500 nucleotides (nt) in length. However, a group of orthologous, metahepadnavirus-derived eHBV elements identified in birds contained some copies that spanned a near-complete genome (**Fig. 2**). Furthermore, in-depth investigation of this insertion demonstrates that it is clearly orthologous across a diverse range of avian species, including eagles, implying that it originated >83 Mya (CI: 77-90 Mya). Even more remarkably, our investigation revealed that an element identified in the tuatara is likely a member of the same group of orthologous insertions. This implies that germline incorporation of the element - labelled *eHBV*-*Meta.1-Sauria* – occurred prior to the divergence of the Lepidosauromorpha and Archosauromorpha ~282 Mya [19]. Given that: (i) we found evidence for independent insertion and fixation of eHBVs at approximately equivalent genomic loci (see above), and; (ii) due to deletion of large regions of terminal eHBV sequence, none of the eHBV-genomic DNA junctions are precisely equivalent on either side of the avian and lepidosaur orthologs, this finding has to be interpreted with caution. However, in each of the pairs of insertions that we propose to have been independently integrated (see **Fig. 4a**), insertions are only located at approximately similar genomic sites. By contrast, the genomic flanks upstream and downstream of the tuatara and avian elements show a strikingly similar arrangement of conserved non-coding sequences (**Fig. 4b**). Since DNA loss is characteristic of Saurian evolution [20] the equivalent genomic region could presumably have been deleted in other major clades descending from the Lepidosaur-Archosaur ancestor.

## DISCUSSION

Our investigation provides a range of new insights into the deep evolutionary history of hepadnaviruses and their impact on animal genomes. Firstly, we show that germline incorporation of hepadnavirus sequences is unique to saurians, despite the fact that hepadnaviruses are known to infect a much broader range of vertebrate groups. It is unclear why germline incorporation is restricted to saurian hosts, but access to germline cells is likely to be a key underlying factor. The relatively high level of genome invasion might be related to specific aspects of transmission and replication in this particular host-virus system (i.e. avi-or herpetohepadnavirus infections in saurian hosts) - particularly as they relate to vertical transmission. Studies of avihepadnavirus infections in domestic ducks show that virus is normally transmitted via vertical transmission *in ovo* and this may be the case in other avian species [21]. Conceivably, herpeto-/avi-hepadnaviruses could have come to rely more on vertical transmission via infection of germline cells than other hepadnaviruses, perhaps in relation to certain aspects of the saurian reproduction system (e.g. internal fertilization and the shelled egg) and the way in which these evolved and this has provided greatly increased opportunity for germline incorporation to occur.

We show that the diversity of hepadnavirus sequences contained within saurian genomes is higher than has previously been appreciated. In particular, the high frequency of germline incorporation in avian lineages allowed an extensive characterisation of ancient avihepadnavirus diversity. Our analysis identified four major subclades within the *Avihepadnavirus* genus, each of which has a relatively broad distribution among avian species. All appear to have circulated throughout a large part, if not most of the Cenozoic Era. However, due to lack genome coverage across avian species, we were only able to obtain an approximate timeline of evolution for each of the four avihepadnaviral lineages. Conceivably, the existence of the four clades might reflect the historical compartmentalisation of avian subpopulations (e.g. due to geographic isolation) during certain periods of their evolution. Currently, we do not have a sufficient level of precision to infer any association between the ancestral distribution of avihepadnavirus strains and the evolutionary history of specific bird lineages. However, the upcoming publication of data from the avian 10K genomes project [22] should allow a much more precise dating of eHBV elements.

We identified several eHBV insertions derived from metahepadnavirus-like viruses, as well as from avi- and herpetohepadnaviruses. These are, to the best of our knowledge, the first metahepadnavirus EVEs to be reported and accordingly they provide a completely new perspective on the evolution of the genus. Remarkably, analysis of the broader genomic landscape surrounding one insertion (*eHBV-Meta.1-Sauria*) indicates that it was inserted into the germline an estimated 280 MYA (CI: 273 - 286 MYA) (**Fig 5a**). This makes *eHBV-Meta.1-Sauria* the oldest EVE described to date, and the first example of an EVE derived from a virus that circulated in the Paleozoic Era (541-252 million years ago).

**Figure 5.**
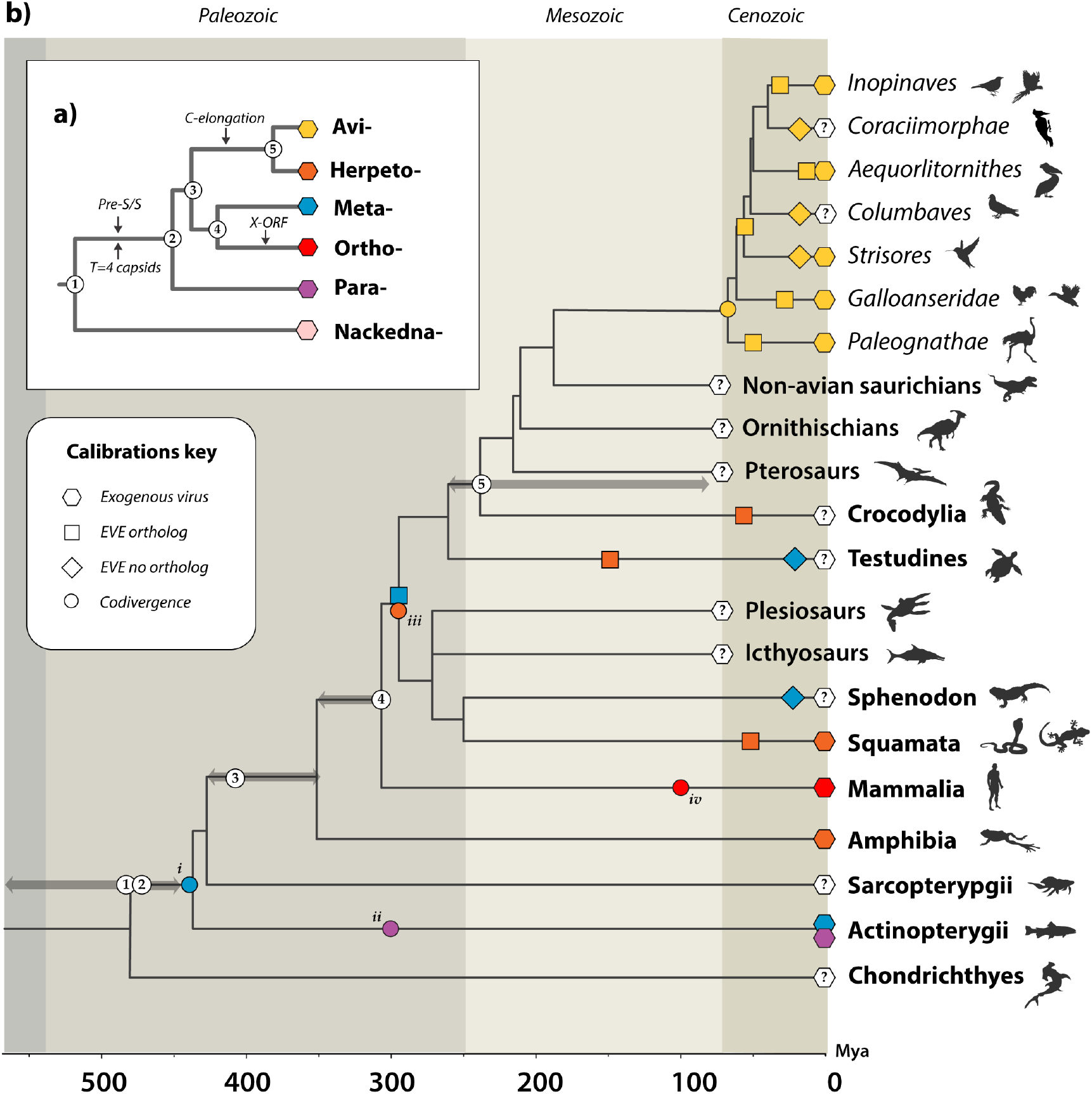
Timeline of hepadnavirus evolution. The inset panel **(a)** shows a schematic phylogeny depicting the established evolutionary relationships between hepadnaviral genera, with black arrows indicating the most parsimonious periods of major evolutionary innovations (after Lauber [4]). Internal nodes are numbered in reference to the time-calibrated phylogeny of vertebrates that is shown in panel **(b**). Panel **(b)** shows a time calibrated vertebrate phylogeny. Geological eras are indicated by background shading. Scale bar shows time in millions of years before present. Colours indicate hepadnaviral genera as shown in panel (a). Shapes on branches indicate four kinds of calibrations as shown in the key. Note that the identification of EVEs that lack orthologous copies (indicated by diamonds) does not allow dates to be inferred, but nonetheless indicates the presence of hepadnavirus in the ancestral members of a given lineage. Numbers in white circles show the putative locations of nodes on the hepadnavirus tree in relation to the timeline of vertebrate evolution. Calibrations based on the assumption of codivergence, as follows; (i) metahepadnaviruses found in fish and saurians; (ii) parahepadnaviruses (found in all teleosts [4]); (iii) herpetohepadnaviruses (assuming that TfHBV originated via interclass transfer from saurians to amphibians); (iv) avihepadnaviruses present in all avian lineages as viruses and/or EVEs. Grey arrows flanking numbered nodes on the host tree indicate time range in which the corresponding virus divergence is estimated to have occurred. Abbreviations: C=Core. S=Surface; EVE=endogenous viral element. Mya=Million years ago.

Metahepadnaviruses have only been identified very recently [5], and along with the parahepadnaviruses (genus Parahepadnavirus) they are the first hepadnaviruses known to infect fish. Whereas the parahepadnaviruses are only distantly related to other hepadnavirus genera, phylogenetic analysis unexpectedly revealed that the *Metahepadnavirus* genus groups as a relatively close sister taxon to the mammalian orthohepadnaviruses in phylogenies [23]. This has been widely interpreted as evidence of cross-species transmission between fish and mammals [3, 5]. However, the discovery that metahepadnaviruses infected ancestral vertebrates challenges these conclusions. We identify metahepadnavirus-derived eHBV sequences derived in a turtle and a lizard (**Table 1**, **Fig. 1**, **Fig. 2**) showing that these viruses not only circulated in saurian ancestors, but also infected the reptile descendants of these organisms. Combined with ancient age of the genus implied by the eHBVs identified in our study, this allows for a simpler explanation of contemporary hepadnavirus distributions, wherein the *Hepadnaviridae* diverge into distinct ‘Meta-Ortho’ and ‘Herpeto-Avi’ lineages prior to the divergence of fish and tetrapods and then subsequently co-diverged (broadly speaking) with their host groups (see **Fig. 5**). As outlined elsewhere, this pattern of evolution does not necessarily preclude zoonotic transmission of related hepadnaviruses (e.g. viruses from the same genus) between related groups of hosts, and phylogenetic analysis does seem to suggest that interclass transmission of herpetohepdnaviruses has occurred between reptiles and amphibians. In general, however, the greater the taxonomic distance between hosts, the less likely a zoonotic jump is to be successful [24].

We show that eHBVs have been intra-genomically amplified in suliforme birds – most likely in association with transposable element (TE) activity - and a large number of these insertions have been fixed. We have previously reported a similar phenomenon for endogenous circoviral elements in carnivore genomes [25], and it has also been described for endogenous retroviruses (ERVs) in primates - for example, hominid genomes contain SVA elements that contain a portion of HERV-K(HML2) [26]. More broadly, it seems that the sequences of certain mammalian apparent LTR retrotransposon (MaLR) lineages, such as the HARLEQUIN elements found in the human genome [27], comprise complex mosaics of ERV fragments. Possibly, the capture of EVE sequences offers a selective advantage to TE lineages. Alternatively, TE sequences containing hepadnavirus-derived DNA might, for some reason, be more likely to be fixed.

Consistent with the idea that germline incorporation of hepadnavirus sequences might, in some cases, be favoured by selection at the level of the host, we identified multiple examples of loci containing multiple fixed eHBV elements, each derived from a distinct germline colonisation event (**Fig 4a**). In principle, the enrichment of eHBVs at specific loci could reflect natural selection – i.e. eHBVs were integrated randomly into genomes, and those integrated at specific loci were selected over time – for example, due to a favourable influence on gene regulation as has been widely reported for TEs and ERVs in animal genomes [28, 29]. However, it could also reflect the preferential integration of hepadnaviruses into these loci (e.g. because they are accessible in embryonic cells).

Comparative studies of eHBVs have been hampered by the challenges associated with analysing these sequences, which are often highly degraded by germline mutation. This may explain why - despite the fact that it has been clear for some time that additional, lineage-specific eHBV insertions are present in some vertebrate species - progress in characterising and analysing novel eHBV sequences has been quite slow. This likely reflects the manifold challenges encountered in identifying and characterising eHBVs. Complicating factors include the hepadnavirus genome structure: the overlapping reading frames and circular genome, both of which can make recovering the ancestral structure of integrated eHBVs less straightforward than it is for other kinds of endogenous viral element. Additional complications arise due to the intra-genomic duplication and re-arrangement of eHBV sequences, and the fact that the hepadnaviral polymerase, which occupies a large proportion of the hepadnavirus genome, shares distant similarity with the reverse transcriptase genes encoded by certain retroelements. While all of these contingencies can be dealt with in one way or another, this is usually done in an *ad hoc* way that makes it difficult for other investigators to recapitulate or build on the work done by previous investigators. In this study we sought to directly address these challenges by using a novel data-orientated approach. This allowed us to publish our findings in the form of an online resource that not only contains all of the data items associated with our investigation (i.e. virus genome sequences, multiple sequence alignments, genome feature annotations, and other sequence-associated data), but also represents the semantic relationships between these data items. Furthermore, via the GLUE engine (a platform-independent software environment) [15] it provides the means to recapitulate all of the analyses performed in our study.

## DECLARATIONS

### Ethics approval and consent to participate

Not applicable

### Consent for publication

Not applicable

### Availability of data and materials

The datasets generated and/or analysed during the current study are <u>publicly available</u> via GitHub.

### Competing interests

The authors declare that they have no competing interests

### Funding

RJG and SL were funded by the Medical Research Council of the United Kingdom (MC_UU_12014/12). GA is funded by ANID FONDECYT 1180705. The funding bodies had no role in the design of the study and collection, analysis, and interpretation of data, or in writing the manuscript.

### Authors’ contributions

Conceptualization, G.A and R.J.G.; methodology, S.L and R.J.G.; validation, S.L. and R.J.G.; formal analysis, S.L. and R.J.G.; writing—original draft preparation, S.L. and R.J.G.; writing— review and editing, S.L. and R.J.G.; visualization, S.L. and R.J.G.; supervision, R.J.G.; project administration, R.J.G.; data curation, R.J.G. All authors have read and agreed to the published version of the manuscript.

## Notes

https://giffordlabcvr.github.io/Hepadnaviridae-GLUE/

### Competing Interest Statement

The authors have declared no competing interest.

