## Supplementary figures and legends for "Ancient evolution of hepadnaviral paleoviruses and their impact on host genomes"

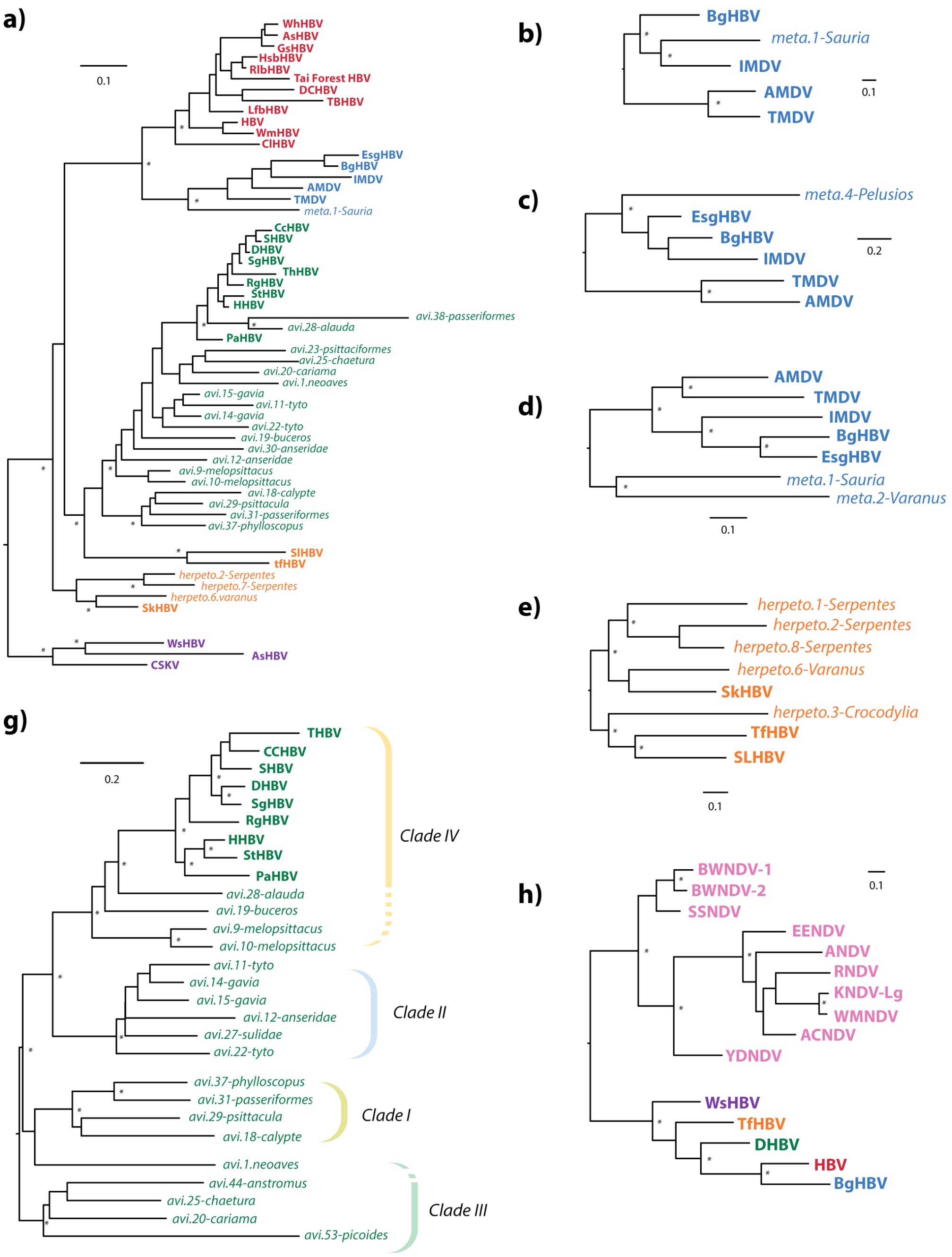


**Figure S1. Evolutionary relationships between eHBVs and contemporary hepadnaviruses. (a)** *Hepadnaviridae* surface (S) protein; **(b)** Metahepadnavirus core (C) protein; **(c)** Metahepadnavirus polymerase (P) protein; **(d)** Metahepadnavirus S protein; **(e)** Herpetohepadnavirus S protein; **(g)** Avihepadnavirus surface (S) protein; **(h)** *Hepadnaviridae* rooted on Nackednaviruses (pink). Asterisks indicate nodes with bootstrap support >70%, based on 1000 replicates. Maximum likelihood phylogenies constructed using RAXML. Virus name abbreviations are as shown in **Table S1**. The dashed bracket for Clade IV denotes that grouping of eHBV elements 9, 19 and 28 does not have high support here, but is supported here but not in all alignment partitions analysed. Asterisks indicate nodes with bootstrap support >=70, based on 1000 replicates. Scale bars show evolutionary distance in substitutions per site.

Colours correspond to viral groups/genera as follows: pink=Nackednavirus; blue=Metahepadnavirus; red=Orthohepadnavirus; orange=Herpetohepadnavirus; green=Avihepadnavirus; purple=Parahepadnavirus.

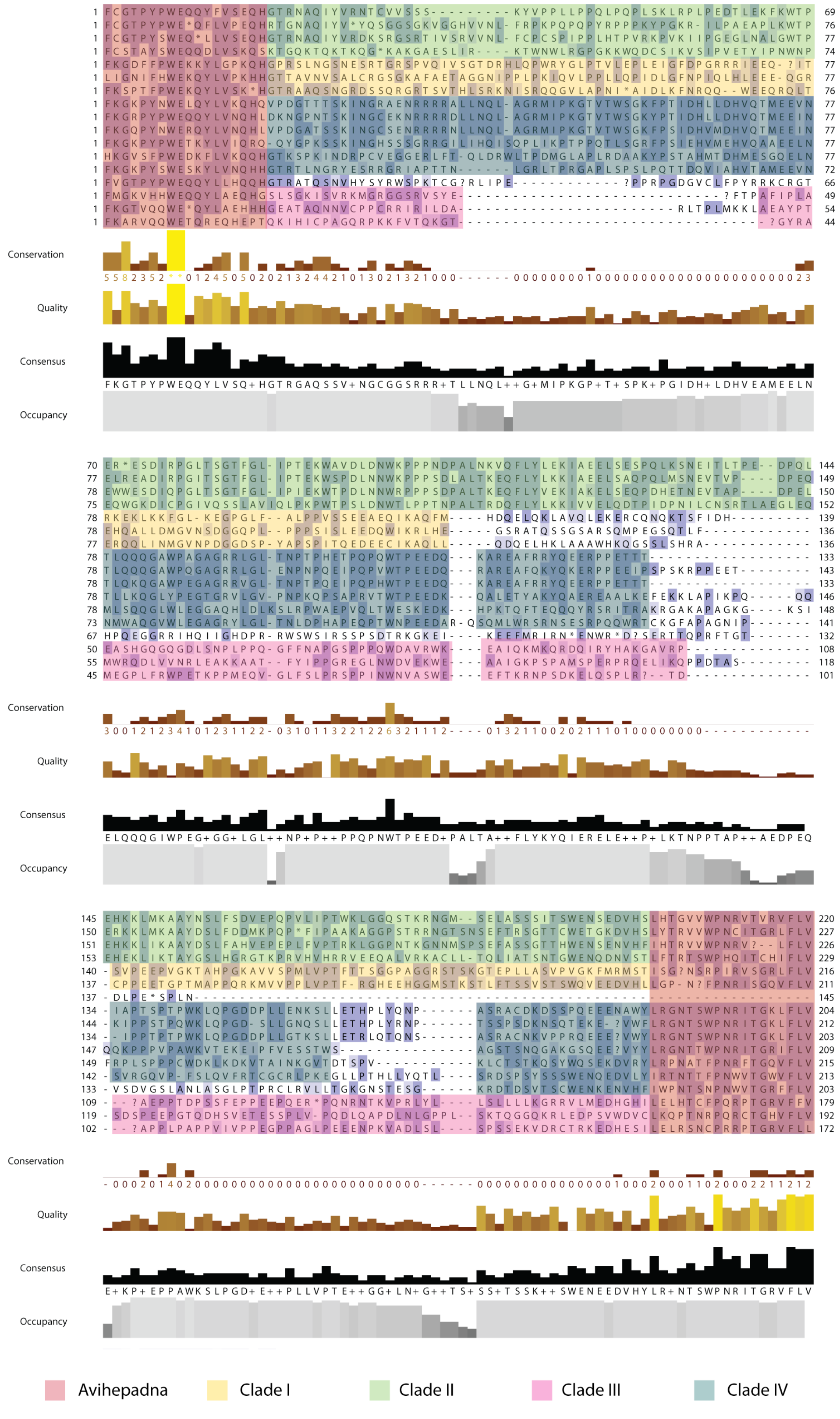


**Figure S2a. Avihepadnavirus clades exhibit distinct variable region 'types'.** The figure shows a section of a multiple sequence alignment of avihepadnavirus protein sequences and translated eHBV elements. The alignment spans variable region 2 in of the avihepadnaviral genome (see **Fig. 2**). Conserved regions of the alignment are shaded darker. Coloured shading is used to highlight region of the alignment. Regions flanking variable region 2, which are conserved across all avihepadnaviruses, are shown in red. Within variable region two there is little obvious conservation of sequence across the entire *Avihepadnavirus* genus. However, conservation can be observed within different members of the individual avihepadnavirus clades and is highlighted by clade-specific colours in line the key.

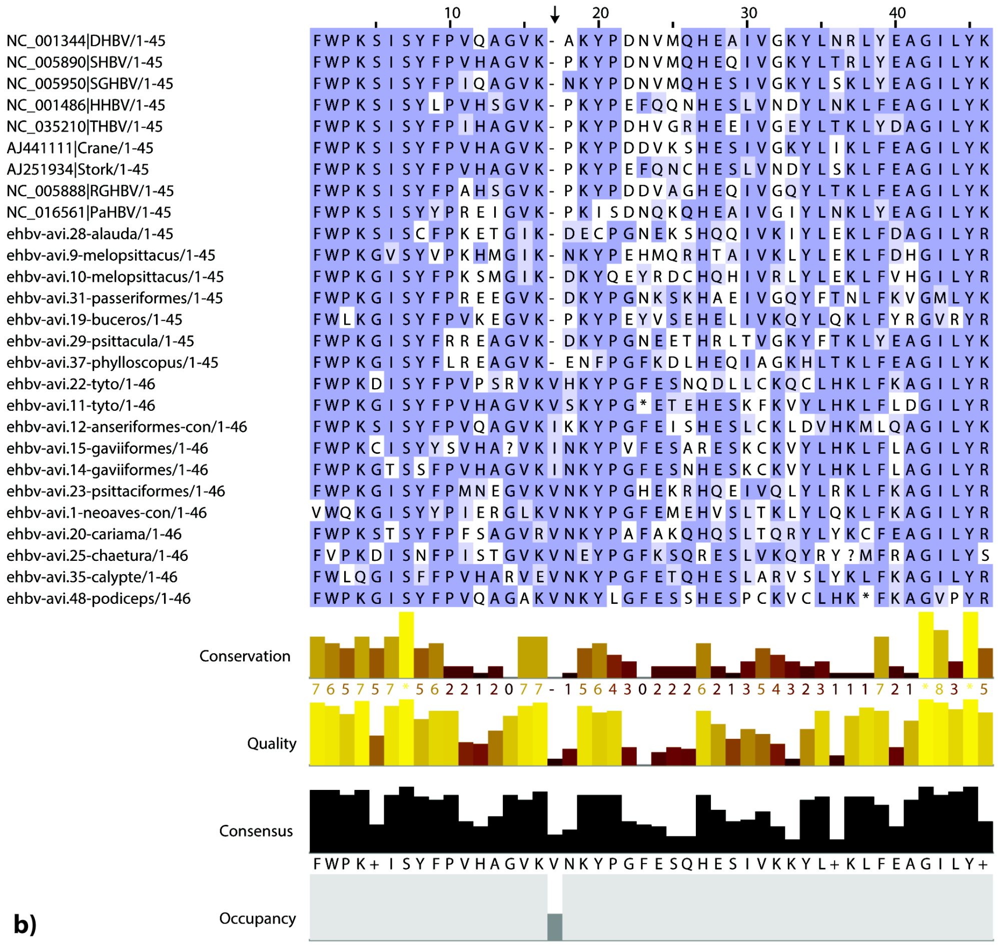


**Figure S2b. A synapomorphic insertion in the avihepadnavirus pol gene. A section of a multiple sequence alignment of avihepadnavirus polymerase proteins.** Taxa names are shown on the left. Black arrow indicates the position of the snyapomorphic insertion K203IVi.

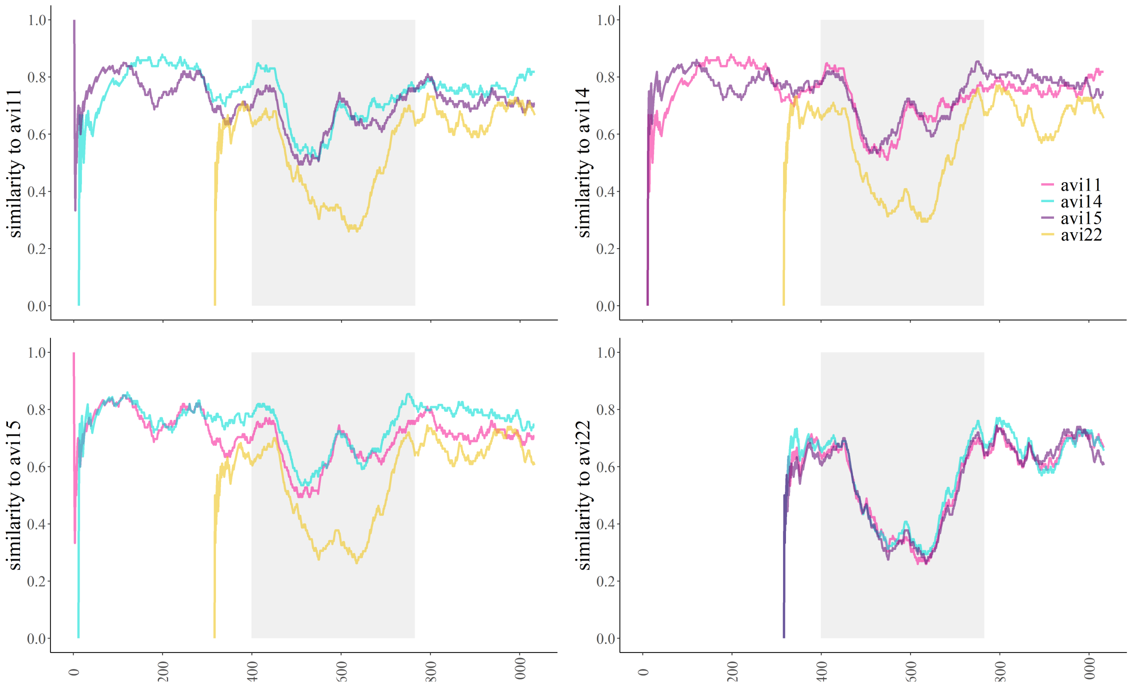


**Figure S3. Pairwise sequence comparisons between selected clade II avihepadnaviruses.** The 4 panels represent comparisons of pairwise similarity of all 3 other clade II avihepadnaviruses against each one of these eHBVs. The x axis corresponds to the alignment's bp positions. Pairwise similarity is plotted at a sliding window size of 100bp. The grey region corresponds to the Pre-S/S (Pre-Surface /Surface) coding region of the genomes.
