## Supplemental table 2 for "Ancient evolution of hepadnaviral paleoviruses and their impact on host genomes"

### Table S2. Previously reported eHBV elements used in this investigation

| Sequence ID (this study) | Previous names | Citation |
|---|---|---|
| ehbv-avi.1-neoaves-con | Zebrafinch EVE, eZHBVc, eAHBV-FRY | [1-3] |
| ehbv-avi.2-estrildidae | eZHBVm | [3] |
| ehbv-avi.3-passeriformes | eZHBV01 | [3] |
| ehbv-avi.4-passeriformes | eZHBV02 | [3] |
| ehbv-avi.5-passeriformes | eZHBV03 | [3] |
| ehbv-avi.6-passeriformes | eZHBV04 | [3] |
| ehbv-avi.7-passeriformes | eZHBV05 | [3] |
| ehbv-avi.8-australiaves | eZHBV0e | [3] |
| ehbv-avi.9-melopsittacus | eBHBV1 | [4] |
| ehbv-avi.10-melopsittacus | eBHBV2 | [4] |
| ehbv-herpeto.1-serpentes-con | eSNHBV1 | [5] |
| ehbv-herpeto.2-serpentes-con | eSNHBV2 | [5] |
| ehbv-herpeto.3-crocodilia | eCRHBV1 | [5] |
| ehbv-herpeto.4-crocodilia | eCRHBV2 | [5] |
| ehbv-herpeto.5-testudines | eTHBV | [5] |
